# How to Scan DNA using Promiscuous Recognition and No Sliding Clamp: A Model for Pioneer Transcription Factors

**DOI:** 10.1101/2023.05.16.541005

**Authors:** Rama Reddy Goluguri, Mourad Sadqi, Suhani Nagpal, Victor Muñoz

## Abstract

DNA scanning proteins slide on the DNA assisted by a clamping interface and uniquely recognize their cognate sequence motif. The transcription factors that control cell fate in eukaryotes must forgo these elements to gain access to both naked DNA and chromatin, so whether or how they scan DNA is unknown. Here we use single-molecule techniques to investigate DNA scanning by the Engrailed homeodomain (enHD) as paradigm of promiscuous recognition and open DNA interaction. We find that enHD scans DNA as fast and extensively as conventional scanners and 10,000,000 fold faster than expected for a continuous promiscuous slide. Our results indicate that such supercharged scanning involves stochastic alternants between local sequence sweeps of ∼85 bp and very rapid deployments to locations ∼500 bp afar. The scanning mechanism of enHD reveals a strategy perfectly suited for the highly complex environments of eukaryotic cells that might be generally used by pioneer transcription factors.

**Teaser:** Eukaryotic transcription factors can efficiently scan DNA using a rather special mechanism based on promiscuous recognition.

## Introduction

The ability to efficiently scan the genomic DNA is an essential feature for all proteins with biological functions that rely on binding to specific DNA target sites (*1*). This requirement applies to most members of the large class of DNA binding proteins (DBP)(*2*), including enzymes involved in DNA repair, synthesis, degradation, editing, and scaffolding. Another important group of DNA scanners are transcription factors (TF), which activate/repress the expression of target genes by locating and binding to cognate sequence motifs present in the relevant control elements (*3*). It is widely accepted that TFs must recognize their cognate motifs specifically in order to perform their function. The specificity in recognizing their cognate motif is supported by a variety of high-throughput selection assays, which have consistently produced well defined sequence binding logos for TFs (*4*). Structurally, specific cognate binding is achieved via detailed interactions formed between the TF and nitrogenous bases from the motif that stabilize the complex in combination with a generic electrostatic attraction to the DNA backbone (*5*). Cognate binding typically results on affinities in the 1nM range, which coincide with the concentrations at which TFs are present in living cells (*6*).

In addition to recognition, finding the target cognate site among hundreds of millions of alternatives that are present in a genome is also a challenge that involves thermodynamic and kinetic considerations (*7–9*). The accepted mechanism for facilitating this search involves an additional non-specific binding mode that recognizes all DNA sequences equally (*10, 11*). Non-specific binding must be weak to avoid outcompeting cognate recognition by sheer numbers (*12*), but it enables a diffusive motion along the DNA that reduces the dimensionality of the stochastic search relative to conventional 3D diffusion-collision kinetics (*11*). A DBP can thus scan the DNA sequence following a spiraling sliding motion around the DNA contour length with diffusion coefficient (*D_1D_*) defined by Schurr’s equation (*13*),

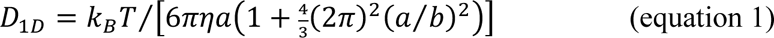

where η is the solvent viscosity, *a* is the radius of the DBP, and *b* is the displacement of a full rotation of the protein around the DNA helix (*b* = 3.4 · 10^−7^ *cm*). Sliding results on full sequence scans, but this motion is significantly slower than linear diffusion due to its rotational component. In addition, equation 1 defines the sliding speed limit, but the actual sliding dynamics should be further slowed by friction. This is so because moving to the next position along the DNA requires breaking, even if transiently, the non-specific interactions that keep the protein bound to the DNA, as well as displacing any loosely associated counterions (*13, 14*). Friction could even be higher *in vivo* due to the abundance of molecular crowders and other DNA associated proteins that can interfere with the sliding motion (*15*). Hence, it has been noted that optimal DNA scanning occurs when 1D and bulk 3D diffusion are mixed (*16, 17*). Mechanistically, maximizing DNA scanning involves balancing the extension of the sliding runs against the friction that ensues from a stronger non-specific DNA association.

1D diffusion on DNA is usually studied using single-molecule fluorescence microscopy (*18*). Such experiments have demonstrated 1D diffusion on many DBPs, including multimeric TFs such as the lac repressor (*19, 20*) and p53 (*21*), restriction enzymes Bam HI (*22*) and EcoRV (*23*), genome editing TALE (*24*) and Cas12a (*25*) proteins, a variety of DNA repair enzymes (*22*), DNA architectural proteins (*26*), and even an adenovirus endopeptidase (*27*). Interestingly, despite the large variety in biological functions, 3D structures, DNA interactions, protein sizes, and even experimental conditions, the existing data reveal a remarkably consistent scanning behavior. For instance, the DBP remains associated to DNA for relatively long times (0.2-10 seconds) in all cases, and moves along the DNA with *D_1D_* values that are within 2.5 orders of magnitude of each other (3·10^−11^–10^−8^ cm^2^ s^−1^). Perhaps more significantly, all of the measured *D_1D_* are just 1-2 orders of magnitude slower than the corresponding sliding speed limits set forth by equation 1, which implies that 1D diffusion generally incurs in little friction, i.e. ≲2.5 *k_B_T* (*22, 26, 28*). The consistency in DNA scanning behavior highlights two major driving factors. One factor involves ensuring that the DNA recognition process is binary, that is, that the non-specific binding mode is uniformly sequence independent. Binary recognition makes the DNA landscape energetically flat for easy sliding. In structural terms, DNA binding becomes sequence independent when it relies exclusively on electrostatic interactions with the phosphate backbone (*29*). The second factor is an interaction interface that encircles the DNA axis to form a sliding clamp (*30*). The sliding clamp can mechanically maintain the DNA association without engaging on strong non-specific interactions, and thus can extend the sliding motion without adding excessive friction. Sliding clamps were first described on ATP-driven DNA polymerases (*31*) and then on DNA repair enzymes (*32*). Importantly, all of the DBPs that have been shown to diffuse on DNA thus far (*19, 21–26*) use binding interfaces that provide some degree of clamping support, whether via tandem arrays of DNA binding motifs/domains or by oligomerizing. The role of the sliding clamp in facilitated diffusion has been carefully examined on TALE proteins, which feature varying numbers of one-base recognition protein motifs that wrap around the DNA axis (*33*), and can diffuse on DNA extensively, even at very high ionic strengths that seriously impair non-specific DNA binding (*24*). An interesting mechanical alternative is a monkey-bar motion that can be performed by proteins that use separate domains for cognate and non-specific recognition connected with a flexible linker (*34*).

However, there is a fundamental group of eukaryotic TFs that control cell fate during embryonic development, morphogenesis, and cell reprogramming (*35*), including homeodomain proteins (*36*), which cannot possibly abide by such DNA scanning rules. These TFs act as pioneers that start global transcription programs by scanning silent chromatin using their ability to recognize cognate motifs in DNA that is both naked or wrapped around nucleosomes (*37*). It has been shown that accessing DNA wrapped on nucleosomes requires that the TF recognizes a cognate motif short enough to be fully displayable on the nucleosome surface (≤8 bp) and uses an open DNA interaction interface to avoid clashes with the histone proteins that comprise the nucleosome core (*38*). Both requirements are in stark conflict with what we understand makes for efficient DNA scanning. A short sequence motif has fewer options to engage in cognate interactions and, hence, is less conducive to binary DNA recognition. In that regard, we recently discovered that the Engrailed homeodomain is actually highly promiscuous, binding DNA with a broad range of affinities that runs proportionally to the similarity of the sequence with its cognate logo (*39*). New high-throughput selection methods specifically designed to sample broad affinity ranges are also starting to report similarly promiscuous profiles for other eukaryotic TFs (*40*). Here is important to note that, by definition, a broad affinity range results on DNA binding landscapes that are energetically rugged, and thus more likely to produce high friction during sliding. Furthermore, the need for an open interaction interface to access DNA on nucleosomes eliminates any clamping support for sliding. The implication is that pioneer TFs must rely exclusively on direct interactions with the DNA to maintain their association while performing 1D diffusion.

The special DNA binding properties of pioneer TFs pose a major puzzle because their biological functions produce even more stringent needs in terms of DNA scanning. As master regulators of cell fate, pioneer TFs control the expression of hundreds of genes(*41, 42*), and operate on DNA regulatory elements consisting of kbp long regions that are localized to nearby a gene (*cis,* intergenic regions) or longer distance along or between chromosome territories (*trans*, enhancers)(*43*). Intriguingly, such long DNA regions contain large clusters of imperfect versions of cognate motifs for key TFs (*12, 44*). These imperfect motif clusters are known to increase local TF occupancy *in vivo* (*45*), and their removal from enhancers impacts cell fate stability during embryonic development (*46, 47*). However, the roles that these motif clusters play in facilitated diffusion remain undefined (*48–50*). A particularly compelling role has emerged in the context of promiscuous DNA recognition, which turns these clusters of imperfect motifs onto a tracking device or transcription antenna that can attract multiple copies of the relevant TFs to co-localize with the region of interest (*39*). Whatever is the role(s) of imperfect-motif clusters in global localization, it is undeniable that such clusters make the DNA landscapes energetically rugged and hence much harder to scan using simple sliding. But there currently is no experimental data available on the facilitated diffusion of pioneer TFs, or on any other protein with comparable DNA binding properties. Therefore, whether or how members of this important group of eukaryotic TFs scan DNA remains unknown.

Here we address this fundamental question by investigating the DNA scanning behavior of the Engrailed homeodomain (enHD) at the single-molecule level. Engrailed is an evolutionary conserved (*51*) master regulator that in *Drosophila* controls cell identity and patterning (*52*). Engrailed controls the expression of over 200 genes, including its own (*53*). In humans, Engrailed-1 is linked to brain and eye defects (*54, 55*) and its misexpression has been linked to cancer (*56*). From a nucleosome targeting standpoint, homeodomains belong to the group of DNA binding domains that can bind DNA all around the nucleosome perimeter (*38*). EnHD is solely responsible for DNA binding in Engrailed, and epitomizes the DNA binding properties of pioneer TFs. The enHD cognate motif is just 6 bp and palindromic (*57*), offering two laterally symmetric target sites. X-ray structures of enHD in complex with cognate DNA demonstrate a wide open interaction interface that lacks clamping support entirely (Fig. 1A). The interface is formed by the lateral insertion of enHD’s C-terminal α-helix (H3) into the DNA major groove in parallel orientation relative to the phosphate backbone (Fig. 1B), which results on interactions with cognate bases through interstitial water molecules, and are hence indirect (*58*). The complex structure points to a strong electrostatic attraction between the two positively charged depressions that flank enHD’s H3 and the DNA phosphate tracks as the major factor stabilizing the complex (Fig. 1A). Furthermore, enHD binds DNA promiscuously both *in vitro* and *in vivo* with sequence preferences that have been integrated onto a statistical mechanical model for predicting the enHD binding free energy landscape of any DNA sequence of interest (*39*). Here we capitalize on such capability to directly compare the 1D diffusive properties of enHD measured by single-molecule fluorescence tracking with this map of its DNA binding energetics. From their vis-à-vis comparison we can uniquely estimate what fraction of the energetic cost of breaking the interactions formed at any given DNA location is converted to friction during DNA scanning. Such detailed information is not generally available for other DBPs, yet it is important to interpret the 1D diffusive process on DNA in mechanistic terms. To examine DNA scanning by enHD we use a variant labeled with the fluorophore A488 at the C-terminal end, which minimizes any interference with DNA binding (Fig. 1B), as we showed previously (*39*). We then employ a dual laser tweezer system to accurately control the position and extension of a single long DNA molecule that serves as scanning substrate, and monitor the 1D diffusive motion of enHD molecules as a function of time and DNA location using correlative confocal fluorescence imaging.

**Fig. 1.**
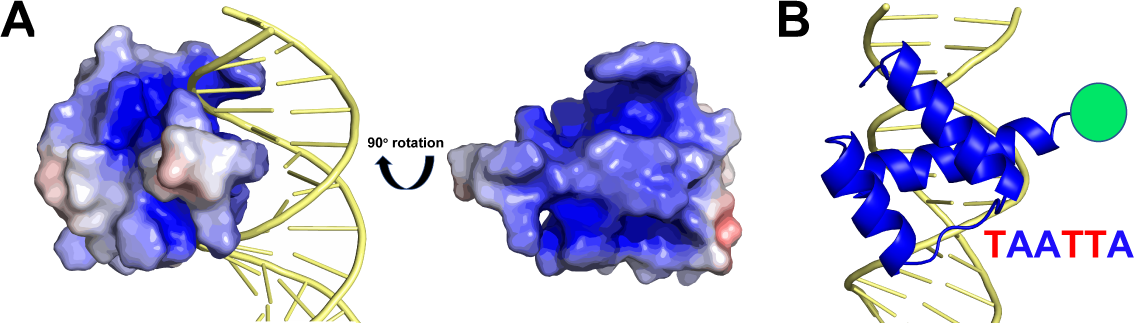
The DNA interaction interface of the Engrailed homeodomain. (**A**) The structure of enHD in complex with DNA (pdb: 1HDD) representing the electrostatic surface of enHD and the DNA in cartoon. The complex shows and interaction interface that is wide open and lacks any DNA clamping effect. The rotated structure of enHD (right) shows the face that directly interacts with DNA, highlighting the strongly positive electrostatic potential of the two depressions that flank α-helix 3 and which interact directly with the DNA phosphate backbone. (**B**) Cartoon representation of the complex illustrating the insertion of α-helix 3 in the DNA major groove parallel to the phosphate backbone. The figure also shows the position of the Alexa 488 fluorophore used for tracking on the DNA as well as the cognate recognition motif for enHD.

## Results

### The genome of phage λ as proxy of an Engrailed control element

As DNA scanning substrate we chose the genome of bacteriophage λ, a 48.5 kbp long DNA molecule that has been widely used as substrate for single-molecule experiments of 1D facilitated diffusion. We examined the binding profile of the full λ-phage genome with the existing statistical mechanical model of enHD promiscuous recognition (*39*). This model, which was parameterized with experimental binding data, calculates the partition function for binding to any of the 2(N-5) sites that are available in any given DNA molecule of N bp. We hence used it to calculate the free energy landscape for enHD binding of the λ-DNA sequence (total of 96,994 possible binding sites) at the same conditions that will be used for the scanning experiments to enable a direct comparison. Fig. 2A shows in teal this free energy landscape converted onto dissociation equilibrium constants integrated over a 45 bp window to reduce site-to-site fluctuations and facilitate visual inspection. The calculation produces a <*K_D_*>_45-bp_ ∼ 4⋅10^−8^ M at 25 mM salt, which confirms that enHD binds on average much more tightly to the λ-DNA than expected for purely non-specific binding. At this resolution the binding profile still shows large local fluctuations in affinity, a few high affinity spikes (∼10^−9^ M) that correspond to 45 bp segments containing two cognate binding sites for enHD (forward and reverse, since the cognate motif is a palindrome), and a central ∼8 kbp region containing clusters of imperfect cognate motifs. Interestingly, the enHD binding pattern of λ-DNA is actually very similar to the profiles we previously reported for the non-coding regions of genes known to be under Engrailed control (*39*), indicating that the λ-DNA is a reasonable proxy of an Engrailed control element. In this light, the central region of λ-DNA could be considered equivalent to a transcription antenna, whereas the spikes recapitulate the random occurrences of cognate sites in any given genome (a frequency of 1 in 4,096 for a 6 bp cognate site). The navy blue profile shows the same equilibrium binding profile integrated over a 320 bp window that is comparable to the position accuracy in our fluorescence tracking experiments (see Methods and Fig. S1). At this resolution the fluctuations in binding affinity are largely averaged out, but the profile still highlights differences in affinity of up to 4-fold along the DNA sequence. From a mechanistic standpoint it is more informative to look at the binding free energy landscape of the λ-DNA at single-site resolution (6 bp). Fig. 2B shows a 1 kbp segment (35.5 to 36.5 kbp) as an example, which illustrates its inherent roughness. This zoomed region shows that the fluctuations in binding free energy between neighboring sites along the λ-DNA genome can be as high as 10 *RT*, resulting in ∼22,000 fold differences in binding occupancy between neighboring sites. The high magnitude and frequency of these free energy fluctuations suggest that enHD should experience extremely high friction when sliding along the λ-DNA sequence.

**Fig. 2.**
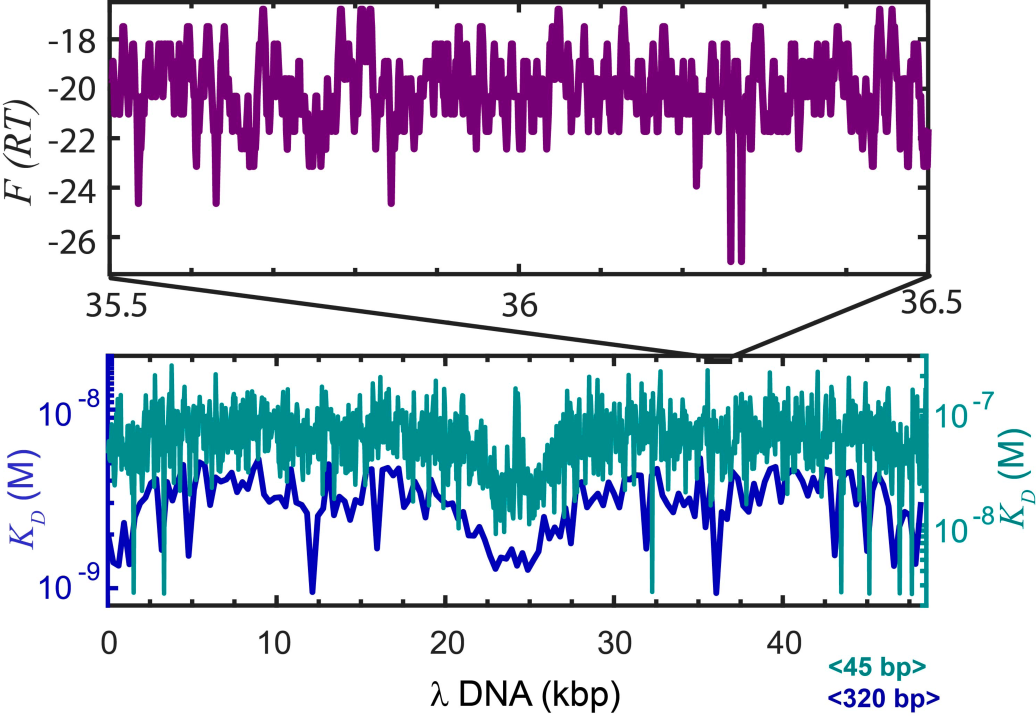
The DNA binding landscape for enHD of the λ-phage genome. The enHD binding profile of the 48.5 kbp sequence of the λ-phage genome calculated with the statistical mechanical model of enHD DNA binding (*39*). The equilibrium dissociation constant for enHD integrated over a window of 45 bp (teal) or 320 bp (navy blue) are shown at the bottom. The top panel shows in purple a 1 kbp detail of the binding free energy landscape (*RT* units) at single-site resolution corresponding to the 35,500 to 36,500 bp segment of the λ-phage genome in the 5’ to 3’direction (binding is bidirectional).

### Optical tweezers and correlative confocal fluorescence microscopy to measure enHD diffusion along the λ-DNA

We use two independent optical traps to mechanically control the position and extension of a single λ-DNA molecule tethered to two beads; and correlative confocal fluorescence microscopy to scan the DNA molecule and track the position of fluorescently labeled protein molecules as they move on the DNA (Fig. 3A). This technique has been recently applied to measure 1D diffusion of Cas12a (*25*). It has the advantage of affording an extremely fine control of the extension and localization of the DNA, which results in a more accurate correlation between the fluorescence signal and the position on the DNA. Moreover, the active mechanical control afforded by the traps eliminates the need of high flows to stretch the DNA, reducing artifacts from hydrodynamic drag. We used λ-DNA biotinylated at both ends to form tethers to streptavidin coated polystyrene beads. The optical traps are used to capture the two beads and hence control the tethered DNA mechanically. Our experimental setup is identical to that used for Cas12a, in which a multichannel microfluidics chip is used to deliver the different components of the assay in sequence (Fig. 3B) using the following workflow: 1) the optical traps are moved to the bottom channel to trap two streptavidin coated polystyrene beads; 2) the trapped beads are moved to the channel with biotinylated DNA to tether the ends of one (and only one) molecule of DNA to both beads using repeated reel-unreel cycles and monitoring tether formation through the force profile of the extension segments; 3) the tethered DNA is stretched to its maximal relaxed extension, and moved to the channel containing fluorophore-labeled enHD; 4) the objective of the confocal microscope is positioned on the DNA and 1D scans are performed in cycle until the DNA molecule detaches, resulting on a kymograph. Fig. 4A shows a 2D image of one molecule of λ-DNA tethered to ∼3.1 μm beads and stretched to its relaxed maximally extended configuration resulting on a separation of 16.5 μm (48,502 x 0.34 nm per base pair). The image also shows several A488-labeled enHD molecules associated to different locations of the λ-DNA. Fig. 4B shows a kymograph constructed from fluorescence confocal line scans along the DNA length, each taking ∼10 ms. The kymograph permits the tracking of single molecules of enHD associated to the λ-DNA molecule as a function of time (x-axis) and position (y-axis).

**Fig. 3.**
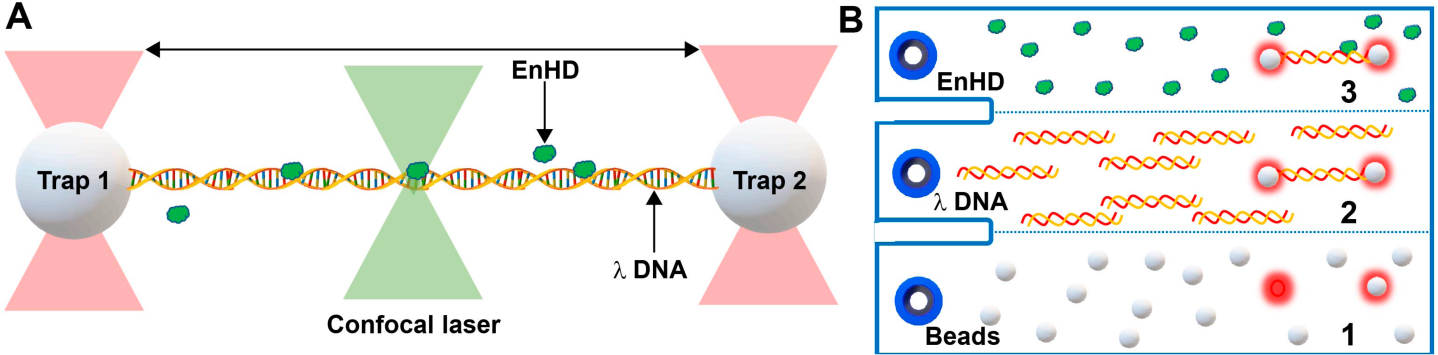
Experimental Setup for Single-Molecule Imaging of DNA scanning by EnHD. (A) Illustration of the experimental setup for imaging the 1D diffusive motion of enHD on the λ-DNA using a dual trap with correlative fluorescence confocal microscope. The two optical traps are used to mechanically control a single copy of biotinylated 48.5 kbp λ-DNA tethered to ∼3.1 micron streptavidin-coated beads. The confocal microscope is scanned along the DNA molecule to image the binding and diffusion of enHD molecules labeled with the fluorophore Alexa 488 at the C-terminus (as shown in Fig. 1B). Line scans were performed in 100 nm steps with 50 μs photon collection time per pixel, resulting on a full scan of the λ-DNA in ∼10 ms. (**B**) Diagram of the microfluidics laminar-flow cell of the instrument showing the workflow for the DNA trapping and single-molecule imaging experiment: 1-streptavidin coated beads are flowed on the bottom channel and the two traps are positioned to trap one bead each; 2-the trapped beads are moved to the middle channel containing the biotinylated λ-DNA at a close distance from one another until one molecule of DNA is tethered to both beads as detected in the force extension profile; 3-the traps with the tethered single DNA copy are moved to channel 3 containing the A488-labeled enHD to perform the scanning studies.

**Fig. 4.**
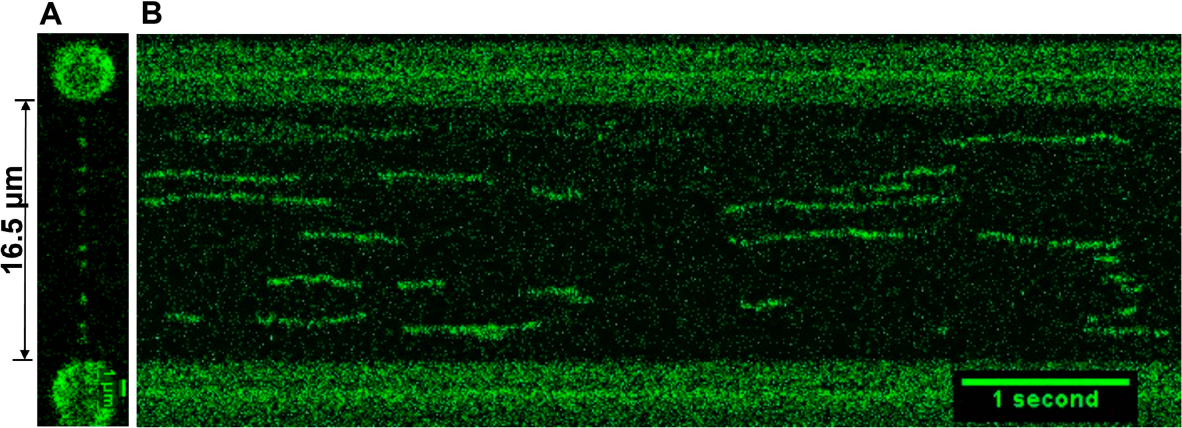
1D Diffusion of EnHD Molecules on the λ-DNA. (**A**) One molecule of the λ-DNA tethered to 3.1 μm beads and mechanically extended to its maximal relaxed extension of 16.5 μm with several A488-enHD molecules bound. (**B**) Kymograph performed with 10 ms line scans along the λ-DNA (y-axis) taken as a function of time (x-axis). The kymograph highlights the dwell time and diffusive motion of individual A488-enHD molecules along the λ-DNA.

### EnHD is an extensive and supercharged DNA scanner

We performed experiments such as those shown in Fig. 4 for enHD wild-type and the mutant Q50K, which enhances the affinity for DNA (*59*). The experiments were performed at various ionic strengths to investigate the effect of modulating DNA binding affinity through the shielding of electrostatic interactions. We used 25 and 50 mM NaCl for the wild-type, and added 75 and 100 mM for the Q50K, taking advantage of its higher DNA affinity. In these experiments we typically obtained several hundreds of trajectories of individual enHD molecules performing 1D diffusion on one λ-DNA molecule. The trajectories were analyzed as described in the methods section to determine the dwell time on the DNA, distance traveled, mean square displacement and average diffusion coefficient for each enHD trajectory. Fig. 5 summarizes the results from such data for the wild-type at 25 mM NaCl (521 trajectories). The data for all other conditions on wild-type and Q50K mutant are given as supplementary information (Figs. S2, S3). Fig. 5A shows a distribution of dwell times that is roughly exponential with a characteristic dwell time on λ-DNA of ∼0.6 seconds. The distribution of traveled distances has a median of ∼1540 bp (or ∼0.51 μm)(Fig. 5B). Therefore, at an ionic strength of 25 mM that is in the middle range used for other facilitated diffusion studies, enHD diffuses along DNA as extensively as do DBPs endowed with DNA interfaces that provide clamping support. Fig. 5C shows the *D_1D_* values for each of the 521 trajectories. These data show a spread of nearly 2.5 orders of magnitude that is again consistent with existing data on other DBPs. The median 1D diffusion coefficient (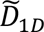) is ∼3.10^−9^ cm^2^.s^−1^, which is in the fast-intermediate range compared to other DBPs. EnHD is a small monomeric protein, and hence its scanning speed is governed by a comparatively fast translational diffusion coefficient. Nevertheless, the 1D diffusion of enHD on DNA is quite fast even in relative terms, as indicated by the fact that is only ∼30-fold slower than its sliding speed calculated with equation 1 (red arrow in Fig. 5C). The slowdown of the enHD 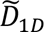 relative to its sliding speed limit is also comparable to those reported on other DBPs.

**Fig. 5.**
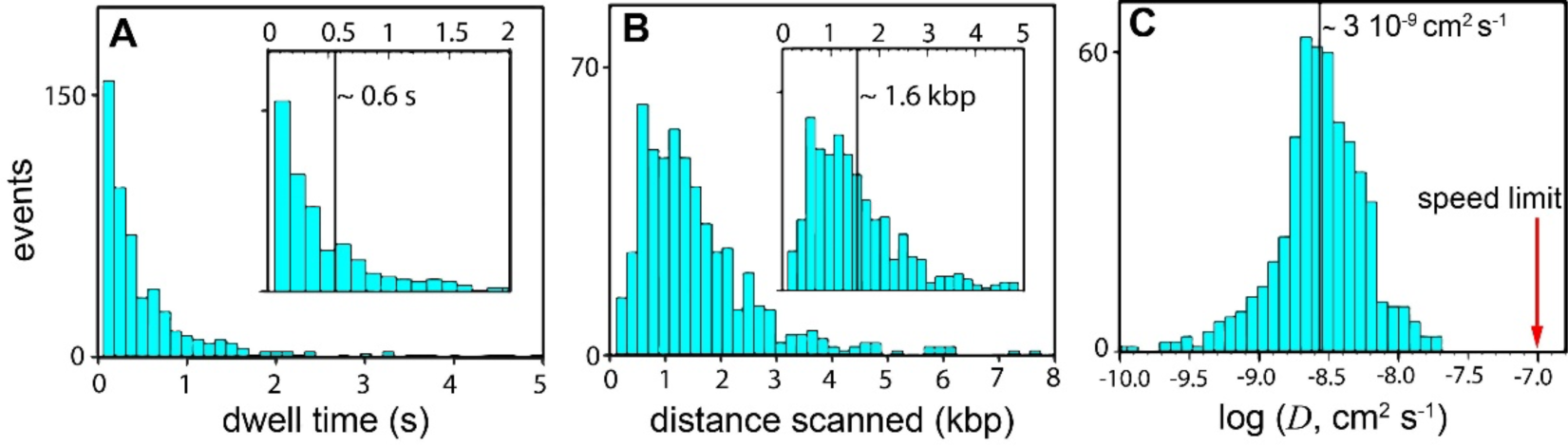
DNA Scanning Properties of EnHD. (**A**) Histogram of dwell times of the wild-type enHD on the λ-DNA. The inset shows the median dwell time as a vertical line on the section of the histogram from 0 to 2 seconds. (**B**) Histogram of the distances scanned along the λ-DNA on single trajectories. The inset shows the median distance scanned as a vertical line on the histogram section from 0 to 5 kbp. (**C**) Histogram of the 1D diffusion coefficients (*D_1D_*) with median indicated as a thick vertical black line. The red arrow signals the sliding speed limit for enHD calculated with equation 1 and *a* =1.7 nm and *b* =3.4 nm.

There is, however, a fundamental difference with other DBPs because enHD is a highly promiscuous DNA binder (*39*). The promiscuous recognition of enHD results on highly rugged DNA binding landscapes, as illustrated in Fig. 2 for the 48.5 kbp genome of the λ-phage. In principle, such landscape ruggedness should proportionally increase the friction that a promiscuous binder encounters during sliding. The question is how much of an acceleration the experimental *D_1D_* of enHD really implies relative to the expectation for a continuous sliding motion on the binding landscape presented by the λ-DNA. Sliding can be described at the microscopic level as a series of discrete steps in which the protein breaks off the interactions that is making with the currently occupied DNA site, moves to the adjacent site, and forms new interactions. Hence, the effective *D_1D_* for sliding results from scaling equation 1 by a friction term 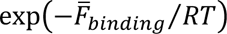 that will depend on the specific DNA binding landscape and experimental conditions. We can estimate this friction term for sliding from the binding profile of the λ-phage genome calculated with our statistical mechanical model (Fig. 2). This calculation indicates that the sliding motion of enHD along the λ-DNA should produce an average friction of ∼21 *RT*. Therefore, the experimental 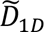 of ∼3.10^−9^ cm^2^.s^−1^ that we have measured for enHD is about 10,000,000 times faster than expected for a continuous sliding motion on the λ-DNA. In other words, the 1D diffusion on DNA of enHD is supercharged relative to sliding, and remarkably impervious to the overall strength and local fluctuations in binding strength that it encounters along the DNA sequence landscape. Such imperviousness to the topography of the DNA binding landscape is further evidenced by the behavior of the Q50K mutant. The Q50K mutation maintains high affinity for the cognate TAATTA sequence, but adds even stronger affinity for the alternate sequence TAATCC (*59*), making its DNA recognition even more promiscuous. To compare the wild-type and the stronger DNA binding Q50K mutant, we introduced an empirical correction for their different affinity based on the ratio between the protein concentrations of the two variants that we had to use in the optical traps-confocal experiments to attain roughly equal binding occupancies on the λ-DNA. The ratio we determined this way is equivalent to a 1.2 *RT* stronger affinity for Q50K. Using this simple correction, we found that Q50K scans DNA with essentially the same 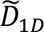 than the wild-type when both proteins are compared at experimental conditions that match their overall affinity for the λ-DNA. These results demonstrate that enHD scans DNA extensively, at rates that are ultrafast relative to a continuous sliding motion, and without being much affected by the ruggedness of the DNA binding landscape.

### EnHD scans DNA with fractional friction relative to its binding free energy

We further explored the level of friction that enHD experiences during DNA scanning by looking at the trends of the aggregated wild-type and Q50K data as a function of ionic strength. Fig. 6A shows the mean and standard deviation of the distribution of stepping rates in ln(bp/s) units for the combined data versus the change in binding affinity calculated by the model. The stepping rate for each trajectory was obtained from the measured *D_1D_* using equation 3 (see Methods), and the data for each condition were then compiled to calculate the mean and standard deviation. The aggregated mean stepping rate data lie on a straight line with negative slope relative to the calculated binding free energy (Pearson coefficient *r* = 0.976). This correlation indicates that: *i*) the effect of the Q50K mutation is indeed reasonably well described by an overall 1.2 *RT* increase in binding affinity; *ii*) ionic strength changes the mean stepping rate (or *D_1D_*) in a manner that is directly proportional to the change in binding affinity calculated by the model. These results support the hypothesis that the diffusion of enHD on DNA is controlled by the breaking and remaking of promiscuous interactions as it moves along the DNA length. However, the slope of the correlation is only −1/3.65 (Fig. 6A). Practically this means that enHD experiences a level of 1D diffusive friction that corresponds to a ∼1/4 fraction of its average free energy of binding to the λ-DNA sequence. There are two scenarios that could explain this result. In one such scenario enHD uses a uniform 1D diffusive motion in which the interactions with DNA are significantly weaker than dictated by the equilibrium binding thermodynamics. The second scenario would involve a hybrid motion in which enHD diffuses along DNA alternating between high friction (DNA bound) and low friction (DNA unbound) modes.

**Fig. 6.**
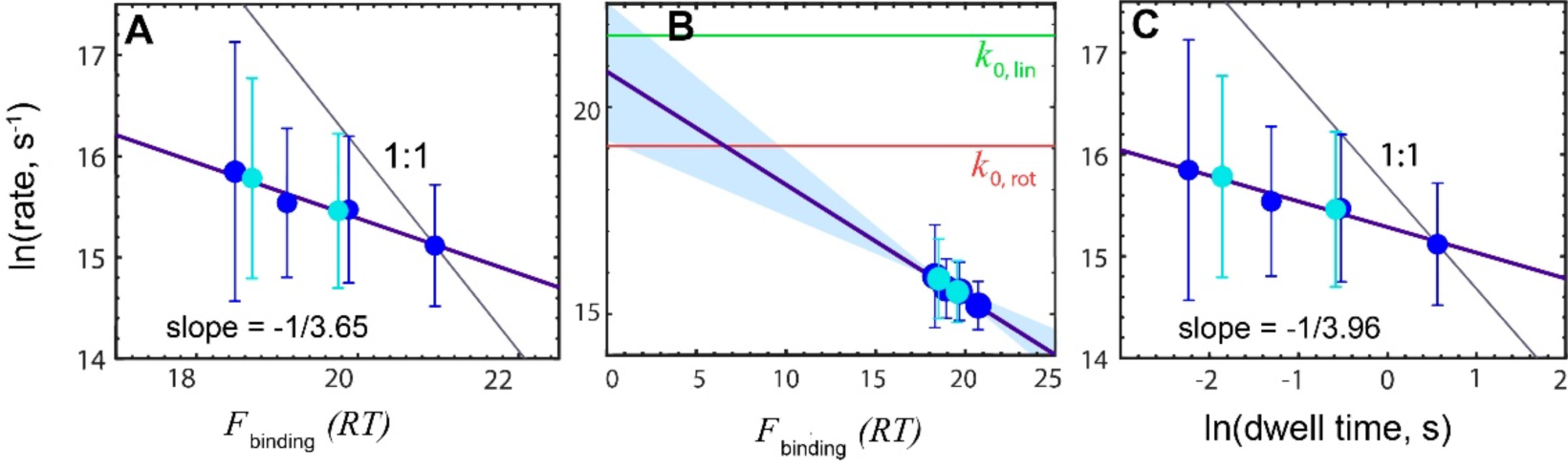
DNA Scanning Speed versus Free Energy of Binding. The *D_1D_* at different NaCl concentrations for the wild-type enHD (cyan) and Q50K mutant (blue) converted to stepping rates in bp.s^−1^ using equation 3, and shown in natural logarithm units, compared to the binding strength in *RT*. The circles indicate the mean and the bars the standard deviation of all the trajectories measured at each condition. (**A**) The experimental stepping rate versus the average binding free energy to the λ-DNA calculated with the statistical mechanical model (*39*). The thick purple line is the linear fit with slope of −1/3.65. The thin black line shows the expectation for a 1 to 1 correspondence. (**B**) As in A but showing the extrapolation of the correlation to zero binding free energy (no friction). The red horizontal line indicates the stepping rate corresponding to the rotational sliding speed limit (as in Fig. 5C) and the green horizontal line the stepping rate for a linear 1D diffusion motion along the DNA with no friction. (**C**) The experimental stepping rate versus the natural logarithm of the experimental mean dwell time on the DNA. The thick purple line is the linear with slope of −1/3.96 and the thin black line shows the trend expected for a 1 to 1 correspondence.

Fig. 6B presents the same data on a scale extended all the way to zero binding free energy, at which point the sliding friction vanishes. The figure also shows the stepping rate limit expected if enHD were diffusing by pure rotational sliding motion without friction (*k_0,rot_*), or performing 1D diffusion with no friction and no rotation (*k_0,lin_*). *k_0,lin_* was calculated from the theoretical translational diffusion coefficient for free enHD with the Stokes-Einstein equation using a hydrodynamic radius for enHD of ∼1.7 nm. *k_0,rot_* was calculated using the same enHD radius and the Schurr equation (equation 1). Here we note that the extrapolation to zero binding free energy reaches an intermediate value between these two limits (Fig. 6B). This suggests that at conditions of ‘zero friction’ the 1D diffusion of enHD on DNA is consistent with a mix of rotational sliding and linear 1D diffusion. Interestingly, the extrapolated rate roughly corresponds to ∼1/4 of the slowdown expected for a pure rotational sliding motion relative to linear 1D diffusion. Therefore, the extrapolated ‘zero-friction’ stepping rate is quantitatively consistent with the fraction of the binding free energy that enHD encounters as scanning friction (i.e. the slope). However, due to the long extrapolation, the confidence interval for this rate is statistically compatible with any value in between the two limits (swath in Fig. 6B).

In Figs. 6A-B we compared enHD’s 1D diffusion with its energetics of binding to λ-DNA as a way to assess the role that promiscuous DNA interactions play on the diffusive process. However, binding is local and site specific, whereas the *D_1D_* of one trajectory is measured over long distances (i.e. ∼1.5 kbp on average, Fig. 5B) and hence reflects averages over hundreds of binding-release events. The implication is that the fractional values for slope (friction) and intercept (dynamics) from the correlation of Figs. 6A-B can still be explained with a fractional friction but uniform scanning scenario. Furthermore, these comparisons are indirect because they rely on calculations with a statistical mechanical model and theoretical estimates of diffusion limits. As an alternative, we directly compared the effects induced by changes in ionic strength on both the stepping rate (*D_1D_*) and dwell time on the DNA. This comparison correlates the rate and extension of 1D diffusion measured from the same trajectories, and should be distinctive for the two scenarios. For instance, if 1D diffusion occurs via a uniform motion with reduced friction, the same interactions that keep the protein on the DNA need to be broken to enable its longitudinal displacement, particularly since enHD binds DNA without clamping it (Fig. 1A). Weakening the electrostatic attraction by ionic screening should affect the stepping rate and dwell time in the same way; that is with a 1:1 correspondence, and both at ∼1/4 of the changes in binding free energy per Figs. 6A-B. In the hybrid scenario enHD alternates between a DNA-bound mode and a low friction mode during which enHD might be partially or fully unbound. Ionic strength will modulate the stepping rate of the DNA bound mode, but should not significantly affect the low friction motion given that enHD is not engaged in interactions with the DNA while in that mode. In such case the interactions formed in the DNA-bound mode still determine the recapture probability after each dissociation-displacement step, and hence how long the protein remains associated to the DNA, or dwell time. These interactions also determine the diffusion coefficient for the DNA-bound mode. However, the difference is that the strength of such interactions will only impact *D_1D_* through the segments of the trajectory during which the protein is actually DNA bound, whereas the low friction segments should be largely insensitive.

Fig. 6C shows the changes in stepping rate (*D_1D_*) versus the changes in dwell time. Consistently with what we observed for the binding free energy (Fig. 6A-B), the changes in dwell time strongly correlate with the changes in stepping rate (Pearson coefficient *r* = 0.983). The linear correlation renders a slope of −1/3.96. That is, the effect of ionic strength on the dwell time is 4-fold stronger than on *D_1D_* (Fig. S4), essentially the same ratio than we observed for the calculated changes in binding free energy. This result is fully consistent with the hybrid DNA scanning mechanism. We note that the observations of Fig. 6 could still be explained with a uniform friction scanning mechanism in the special case in which the ionic strength has two counterbalancing effects on 1D diffusion: i) weakening of the enHD-DNA interactions, which would reduce the dwell time and increase the stepping rate (*D_1D_*) proportionally; and ii) a reduction of the stepping rate by added friction resulting from the forced displacement of an increasing number of counterions associated to the DNA (*13*). However, the comparison between wild-type and Q50K at the same ionic strength shows that the decrease in stepping rate (*D_1D_*) of the Q50K relative to the wild-type is again 1/4 of the 1.2 *RT* enhancement in binding affinity caused by the mutation. In other words, enHD experiences a small fraction of the free energy it uses to interact with the DNA as scanning friction, regardless of whether the binding strength is tuned by ionic strength or mutation, indicating that such effect is not caused by counterion-induced friction. Overall, these results strongly support a hybrid mechanism for DNA scanning in which enHD alternates between high and low friction modes.

### Large heterogeneity in 1D diffusive behavior

We also noted that the enHD’s *D_1D_* was highly variable among individual trajectories (Fig. 5C). The resulting variations in scanned distance are quite significant, as illustrated by the two trajectories of wild-type enHD in Fig. 7A, which are of comparable duration and take place in nearby locations on the λ-DNA but result in largely different traveled distances. We looked closer into such heterogeneity by calculating *D_1D_* over short time intervals along each 1D diffusive trajectory and hence obtain a distribution of ‘quasi-instantaneous’ *D_1D_* values. This approach has been used before to analyze the interconversions between search (diffusing) and recognition (static) modes that occur in timescales close to 1 s for some DNA repair enzymes (*60*). The distribution of ‘quasi-instantaneous’ *D_1D_* was distinctly bimodal in that case, which permitted to resolve the 10-30 fold differences in the characteristic 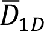 of the two modes (*60*). A similar analysis of our enHD data rendered several thousands of *D_1D_* values per experimental condition. Fig. 7B shows the distribution for the ∼9,000 *D_1D_* values of the Q50K mutant at 50 mM NaCl, which we use here for illustration because these data give the best compromise between number of measured trajectories and their duration. However, the same trends are present in all of the other experimental conditions, including the wild-type at 25 mM salt (Fig. S2).

**Fig. 7.**
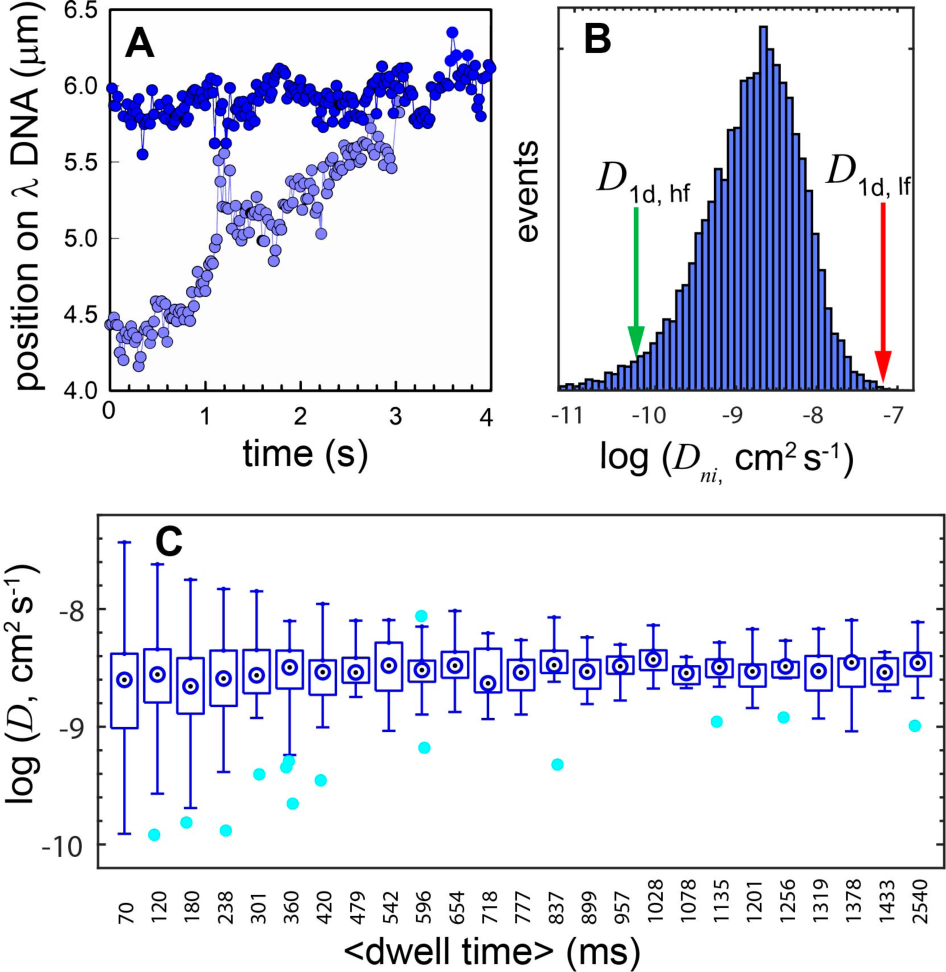
Heterogeneous DNA Scanning by EnHD. 1D diffusive properties on the λ-DNA of the Q50K mutant of enHD at 50 mM NaCl. (**A**) Exemplary diffusive trajectories as a function of time with largely different displacements occurring on nearby locations of the λ-DNA are shown in different shades of blue. (**B**) Histogram of ‘nearly-instantaneous’ *D_1D_* obtained by binning all the 1D diffusive trajectories in 30 ms intervals. The characteristic *D_1D_* values considered for the high-friction and low-friction modes are shown as green and red arrows, respectively. (**C**) Box plot showing the variation in overall trajectory *D_1D_* as a function of the duration of the scanning trajectory on the λ-DNA. Whiskers show the end points, box edges the lower-upper quartiles, and dotted circles the bin medians. Cyan circles are outliers.

For enHD, we find a distribution of ‘quasi-instantaneous’ *D_1D_* values that extends over 3-orders of magnitude, indicating that its 1D diffusion is vastly heterogeneous (Fig. 7A). The distribution is also broadly unimodal in a log_10_ scale, with maximum at ∼2·10^−9^ cm^2^s^−1^ for Q50K at 50 mM NaCl. A unimodal distribution could still reflect a single mode that happens to be severely broadened by experimental error. In these experiments the error originates from the position accuracy of our measurements (∼105 nm, see Methods and Fig. S1) and the short time intervals that we used to determine *D_1D_* (30 ms). Using simple error analysis, we estimate that both factors combined propagate to an experimental uncertainty of ± 0.8⋅10^−9^ cm^2^ s^−1^ in *D_1D_*, which would result on a total data spread (i.e. 4 standard deviations) of ∼0.5 orders of magnitude. The variance in ‘quasi-instantaneous’ *D_1D_* that we observe for enHD is thus many orders of magnitude larger than the experimental error. Another important source of variability in the ‘quasi-instantaneous’ *D_1D_* is the local topography of the DNA binding landscape. We can estimate this factor using the enHD binding landscape of λ-DNA from the statistical mechanical model. We thus calculated the λ-DNA binding profile with a rolling average of 320 bp that is comparable to the position accuracy of our measurements (Fig. 2, navy blue), which indicates that the mean binding affinity between neighboring 300 bp segments of the λ-DNA changes by ≤ 5-fold. Such landscape roughness combined with the experimental error mentioned above propagate to an expected total spread in *D_1D_* of ∼0.8 orders of magnitude, which is still far too little to account for the vast heterogeneity present in the data (Fig. 7B).

In fact, the heterogeneity in ‘quasi-instantaneous’ *D_1D_* for enHD seems covers the full range that is either physically plausible or measurable. The fastest *D_1D_* values are very close to the sliding speed limit, but do not bypass it (red arrow in Fig. 7B). As for the other extreme case, 5·10^−11^ cm^2^s^−1^ represents the lowest value that we can practically determine with minimal accuracy given the short displacements that occur in 30 ms when the diffusion coefficient is below this limit. Hence, the fact that the log_10_ distribution is unimodal despite the massive spread of >3-orders of magnitude suggests that such heterogeneity might be caused by dynamic averaging between scanning modes with vastly different *D_1D_* and which interconvert in timescales comparable to the 30 ms time window. We further investigated this issue by looking at the *D_1D_* variance as a function of the duration of the trajectory. We classified all the 1D trajectories from a given condition into bins according to their dwell times, and performed a simple statistical analysis of the data within each bin. Fig. 7C shows a box plot that summarizes the results of such analysis for Q50K at 50 mM (results for the wild-type at 25 mM salt in Fig. S6). The plot reveals that the full spread in *D_1D_* from full trajectories (indicated by the whisker ends) is inversely proportional to the dwell time. The spread spans 2.5 orders of magnitude for the bin containing the shortest trajectories (∼70 ms), and steadily decreases for longer trajectories, eventually leveling off at ∼0.5 orders of magnitude. A basal level of 0.5 orders of magnitude is in fact consistent with our estimated experimental error (see above), which is reasonable considering that a *D_1D_* measured over a 0.8 s trajectory reflects averages of ∼30 samples from the ‘quasi-instantaneous’ distribution of Fig. 7B, and of over 2.2 kbp displacements along the λ-DNA. On the other hand, the 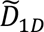 per bin is essentially constant: (2 ± 0.36).10^−9^ cm^2^.s^−1^, indicating that the data contained within each bin provides reasonable sampling of the underlying distribution, with perhaps the exception of the longest time bins containing few datapoints. The trend in the binned data indicates that the heterogeneity in *D_1D_* from trajectory to trajectory reflects a partial dynamic averaging of inherently distinct diffusive behaviors. The detected pattern is strongly suggestive of a stochastic process in which enHD alternates in tens of ms timescales between scanning modes with vastly different *D_1D_*.

### Signatures of the hybrid DNA scanning mechanism for enHD

The simplest mechanism that can explain all of our enHD 1D diffusion data is a DNA scanning process that involves stochastic alternants between high-friction (slow) and low-friction (near the speed limit) modes. We explored the implications of such scenario by performing stochastic kinetic simulations with an elementary model in which enHD interconverts between two scanning modes with *D_1D_* defined by the limits shown in Fig. 7B: a low-friction mode with *D_1D,lf_* = 6⋅10^−8^ cm^2^ s^−1^ (red arrow) that is consistent with the sliding speed limit, and a high friction mode with *D_1D,hf_* = 5⋅10^−11^ cm^2^ s^−1^(green arrow). The latter is likely an upper bound because we do not have sufficient resolution to accurately determine ‘quasi-instantaneous’ *D_1D_* values below 10^−10^ cm^2^ s^−1^. The model assumes that enHD performs 1D diffusion by stochastically alternating between the two modes with exponentially distributed times according to mean values 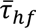 and 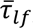, defined such that 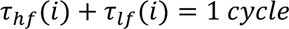 and 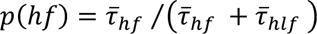 (see Methods). Using switching cycles as the basic unit, the model can be fully determined from the ratio between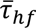 and 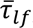 as only parameter, which we adjusted to reproduce the overall trends of the enHD experimental data. The results from these stochastic simulations confirm that such a hybrid high-low friction scanning process results on unimodal distributions of observed *D_1D_* (in log_10_ scale) with variance that is heavily dependent on the number of switching cycles that occur during the trajectory (Fig. 8A). For instance, when the trajectory is shorter than one switching cycle, the observed *D_1D_* values cover the full range defined by the high- and low-friction extremes. The distribution narrows down significantly after two cycles, and the variance keeps on decaying for increasing numbers of cycles, as anticipated. Despite the elementary nature of this model, the simulations closely recapitulate what we observe in the experiments (Fig. 7B-C) using physically plausible parameters. Such agreement provides strong support to the hypothesis that enHD stochastically alternates between high and low friction 1D diffusive modes.

**Fig. 8.**
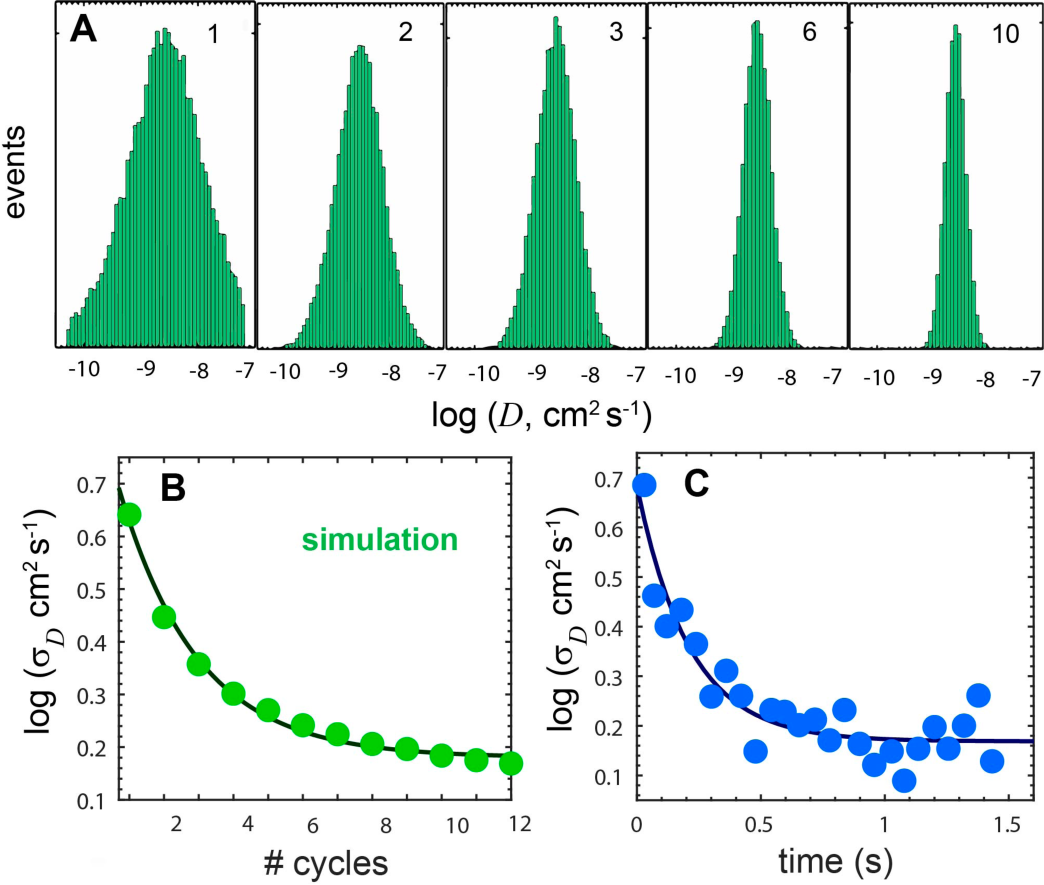
Signatures of an Alternating High-Low Friction Scanning Mechanism. The characteristics of the hybrid motion that involves stochastic alternants between a high-friction and a low-friction mode can be inferred from an analysis of the *D_1D_* variance. (**A**) Examples of histograms of the distribution of *D_1D_* per trajectory for different numbers of switching cycles occurring in the trajectory (1, 2, 3, 6, 10) obtained with stochastic kinetic simulations of the high-low friction switching model described in the text. The simulations were performed with the characteristic *D_1D,hf_* and *D_1D,lf_* values from Fig. 7C, *p(hf)* = 0.96 and *p(lf)* = 0.04. (**B**) Changes in *D_1D_* variance (in log_10_ units) as a function of the number of cycles (1 to 12) obtained from the same stochastic kinetic simulations. The curve represents a fit to an exponential decay function with relaxation constant of ∼2.2 cycles and an offset of 0.185. (**C**) Experimental changes in *D_1D_* variance (in log_10_ units) as a function of trajectory duration for the Q50K mutant at 50 mM NaCl data from Fig. 7C. The curve represents a fit to an exponential decay with relaxation time of ∼210 ms (95% confidence interval of 150-340 ms) and offset of ∼0.17 (95% confidence interval of 0.137-0.204).

The simulations also provide a useful framework for estimating the switching frequency from the experimental *D_1D_* variance. Fig. 8B shows a plot of the standard deviation of *D_1D_* (in log_10_ units) as a function of the number of cycles derived from the simulations summarized in Fig. 8A. The analysis of variance for such data is complex, but we found that the log(*σ*_*D*1*D*_) can be reasonably approximated to an exponential decay with a rate of ∼1/(2.2 cycles) plus an offset (see fitted curve in Fig. 8B). The offset is just an empirical correction to account for the slow asymptotic decay to zero as the number of cycles tends to infinity. Fig. 8C shows the same analysis for the experimental data of Figs. 7B-C as a function of time, which has very similar trends in the magnitude of the log(*σ*_*D*1*D*_) and its decay as a function of time. Curve fitting of the experimental data in Fig. 8C indicates that the decay in variance can be approximated with a relaxation time of ∼210 ms. We convert this relaxation time to cycles using the simulation data as reference, which leads to an estimated high-low-friction cycle of 210/2.2 = ∼95 ms. Using the 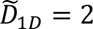·10^−9^ *cm*^2^*s*^−1^ we determined at these experimental conditions (for Q50K at 50 mM NaCl) we then estimate that enHD travels an average of ∼195 nm (∼600 bp) per 95 ms cycle consisting of one low and one high friction step. The basal level in experimental *D_1D_* variance estimated from the curve offset parameter should contain contributions from the still finite number of switching cycles contained in the longest trajectories together with contributions from experimental error and landscape ruggedness. The fitted offset was σ = 0.9⋅10^−9^ cm^2^ s^−1^, which corresponds to a maximal spread (4σ for 95%) of ∼0.78 orders of magnitude. This spread is consistent with the error propagation analysis discussed in the previous section. Therefore, the analysis of *D_1D_* variance is quantitatively consistent with a hybrid high-low friction scanning process in which enHD undergoes a limited number of stochastic interconversions per diffusive trajectory. We further note that this scenario also explains the increased spread in *D_1D_* that occurs at higher ionic strength (whiskers in Fig. 6) as being a direct consequence of the shorter dwell times on the DNA, and hence reduced dynamic averaging, that result from raising the ionic strength (Figs. S2, S3).

The high-low friction hybrid scanning model allows to make other useful inferences about the mechanism underlying the experimental data. Because the *D_1D_* for the high- and low-friction modes define the scanning limits that are accessible to enHD, the effective *D_1D_* of any given trajectory represents how much time/distance enHD spends/travels using each mode. In other words, the 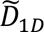 contains specific information about the static probabilities for the high- and low-friction scanning modes. In this respect, the more the two modes differ in *D_1D_* the stronger is the net effect that a rare mode has on the 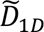. With the low and high friction *D_1D_* values indicated in Fig. 7B, and given 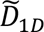 = 2 · 10^−9^ *cm*^2^*s*^−1^, we can thus estimate that enHD (Q50K mutant at 50 mM) spends most of the time during 1D diffusion engaged in the high-friction DNA-bound mode (p ∼0.97), and only 3% in the low-friction mode. Remarkably, the trends are reversed in terms of scanned distance, and hence enHD covers ∼84% of the total traveled distance during the rare, but also so much faster, low friction scanning segments. These estimates are equivalent to mean scanning displacements per cycle of ∼27 and ∼170 nm (∼80 and ∼520 bp) for the high- and low-friction modes, respectively. For the wild-type at 25 mM the numbers for the low-friction mode change slightly: *p*∼0.045 and 86% traveled distance, owing to the slightly faster 1D diffusion that occurs in this variant (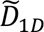 = 3· 10^−9^ *cm*^2^*s*^−1^, Fig. 5C).

## Discussion

### Physical determinants for DNA scanning without clamping support

Here we show that enHD carries out 1D diffusion on DNA that is extensive and fast relative to its theoretical speed limit (Fig. 5), and on par with oligomeric multidomain TFs with complex DNA interfaces, such as p53 (*61*). A key difference relative to previously studied DBPs is that enHD binds DNA with a fully open interaction interface that engages the DNA laterally (Fig. 1A). Such an interface can only maintain the mechanical association of the protein by engaging in direct interactions. Our results on enHD thus demonstrate that a sliding clamp is not a necessary requirement for achieving extensive DNA scanning. The unexpected DNA diffusive properties of enHD shed new light on the structural and energetic factors that act as drivers of facilitated diffusion by DBPs in general, and especially by those that lack clamping support. The structure of enHD in complex with DNA points to a strong electrostatic attraction between the protein and the DNA phosphate backbone as the key energetic factor holding them together during 1D diffusion. EnHD’s electrostatic potential is indeed strongly positive due to an accumulation of positive charge on the face that directly interacts with DNA (Fig. 1A) that is known to be highly destabilizing of the enHD folded structure (*62*). Recent work has shown that such charge distribution enables an electrostatic spring-loaded mechanism that enHD uses for the conformational control of cognate DNA recognition (*63*). From a DNA scanning viewpoint, the electrostatic attraction may play a special role relative to specific DNA interactions because of its action at longer distances. The distance range for electrostatic interactions in aqueous solution is determined by the Debye length. While the actual Debye length for protein-DNA interactions remains undetermined, theoretical arguments indicate that at the very least is 1 nm (*9*). A Debye length of 1 nm, or possibly longer, could perhaps be sufficient to maintain a loose but continuous electrostatic association between enHD and DNA over the entire 1D diffusive trajectory that we observe, including the superfast low-friction mode.

In general, our experimental results provide strong support for a long range electrostatic attraction as the dominant factor leading to enHD’s extensive 1D diffusion on DNA. Namely, we find that moderate increases in electrostatic screening by ionic strength reduce the dwell time of enHD on DNA drastically. For example, raising the salt concentration from 25 to 100 mM decreases the dwell time of Q50K by 20-fold (3 natural log units, Figs. 6C, S4). In fact, the ionic strength affects the dwell time of enHD on DNA in the same proportion than it does its thermodynamic binding affinity, as demonstrated by a slope of ∼0.9 in the correlation between predicted binding affinity in *RT* units and the experimental ln(dwell time) (Fig. S5). This result implies that the electrostatic interactions that control 1D diffusion are of the same magnitude than the electrostatic contributions to enHD’s DNA binding affinity. The Q50K mutant provides further evidence that is in this case independent of the ionic strength. Q50K has one extra positive charge located right on the interface with DNA that enhances the overall DNA binding affinity (*59*). Correspondingly, at the same ionic strength Q50K dwells on the DNA longer than the wild-type in the exact same proportion than its binding affinity increases (Fig. S5). Taken all of these together, we conclude that what makes enHD hold onto DNA for such long times during 1D diffusion is primarily the strength and effective range (Debye length) of the attractive electrostatic potential between them.

Further inspection of the enHD-DNA complex structure reveals some structural features that may facilitate an extensive diffusive motion along the DNA that is driven entirely by an electrostatic attraction. Particularly, H3, which protrudes out of the enHD core, is flanked by two depressions that accumulate most of the positive electrostatic potential of the protein. In the complex with DNA, the insertion of H3 into the major groove interlocks these positively charged depressions with the two tracks of the phosphate DNA backbone resembling one half of a zipper slider (Fig. 1A). This one-sided zipper slider interface might be essential to keep the protein onto the DNA tracks via a long-range electrostatic attraction in the absence of promiscuous interactions with the bases.

### Hybrid DNA scanning as strategy for navigating rugged binding landscapes

Facilitated diffusion assumes a binary interplay between cognate binding, used for recognition, and weak non-specific binding for DNA scanning (*11*). These elements generally result on a search landscape that is one-dimensional and flat, with a single minimum at the target site. However, enHD’s promiscuous recognition drastically changes the DNA scanning scenario since the sequence landscape becomes highly rugged (e.g. Fig. 2), which should impair 1D diffusion and make sliding extremely sluggish. Yet, our results show that enHD diffuses along DNA extensively and with a median *D_1D_* that is close to the fastest previously characterized DBPs, and only ∼30-fold below its theoretical speed limit (Fig. 5C). This is a striking result that implies that enHD scans DNA about 10,000,000 times faster than it would be expected from a continuous sliding motion. The DNA scanning mechanism of enHD is thus supercharged and impervious to the topography of the sequence binding landscape. We find that at the core of such scanning behavior is a hybrid 1D diffusive motion composed of stochastic alternants between high-friction and low-friction modes. In the high-friction mode enHD sweeps the local DNA sequence, whereas the low-friction mode allows enHD to deploy to another location to perform the next local sweep. This mechanism has some parallels with the way enzymatic DBPs search for their targets by switching between an active scanning (search) mode and a recognition state that locks the enzyme at the target site to perform the catalysis. The inter-mode conversions in these enzymes are usually slow and can be controlled externally, or even stalled, via the catalytic step. This feature was instrumental to resolve both modes in single-molecule trajectories (*25, 64*) or from a distribution of ‘nearly-instantaneous’ *D_1D_* values (*60*). The hybrid 1D diffusion performed by enHD is different, however, because it only involves the scanning function (no catalysis) and is purely stochastic and independent of target recognition. The enHD switching between high- and low-friction modes is too transient to be fully resolved in single-molecule diffusive trajectories, but it does become evident in the stochastic analysis of the single-molecule data. The analytical approach that we use here for that purpose is similar to the statistical optimal estimator procedure that Vestergaard et al. applied to the 1D diffusion on DNA of the enzyme hOggI (*65*). The statistical analysis is obviously indirect, but still sheds important light on how enHD avoids getting trapped in the myriad local minima that it encounters in the binding landscape of any natural DNA sequence. The hybrid scanning mechanism we identify on enHD could be potentially used by any other promiscuous DNA binder, including pioneer TFs which must also forego the sliding clamp for functional reasons.

For instance, we find that enHD spends most of the scanning time in the high-friction mode (*p* > 95%), which is when the DNA sequence is actively probed. High-friction steps last about 90 ms on average, and result on mean DNA displacements of ∼27.5 nm (∼85 bp), or ∼1 bp per ms. This rate is actually comparable to the *k_off_* for release from the cognate site that we have recently measured for enHD using single-molecule Förster resonance energy transfer methods at equivalent experimental conditions (unpublished results). Therefore, the time that enHD spends engaged in the high-friction mode appears sufficient to fully probe the local DNA sequence via a series of promiscuous binding-release events. The much rarer low-friction mode plays a complementary role that enables enHD to quickly deploy to alternative neighboring regions and resume scanning. Our analysis indicates that the low-friction mode consists of ∼3-4 ms bursts that occur stochastically every 95 ms, or ∼10 times per second. During a low-friction burst enHD diffuses an average of ∼173 nm along the DNA (∼520 bp), which means that this mode is responsible for 80% of the total distance traveled along the DNA. The reason for these numbers is that the low friction mode is extremely fast, even faster than the rotational sliding speed limit according to the extrapolation of the experimental stepping rate (*D_1D_*) to ‘zero friction’ conditions (Fig. 6B).

An important issue that follows is whether the properties we estimate for the low-friction mode are actually consistent with the long diffusive trajectories detected by fluorescence single-molecule tracking (e.g. Fig. 7A). 3-4 ms is indeed a shorter interval than the time resolution of our correlative optical tweezers confocal microscope, which takes ∼10 ms to scan the full λ-DNA. The lateral optical resolution of our confocal microscope is ∼240 nm, and the position accuracy after point-spread function analysis is ∼105 nm (see Methods and Fig. S1), which are comparable to the estimated mean displacement in the low-friction mode. Our experimental resolution is thus insufficient to resolve the low-friction bursts, but it should be capable of revealing their footprints in the measured trajectories. For instance, the rare transitions to the low-friction mode should typically occur while the confocal microscope is scanning at a distant region in the λ-DNA, and hence they will be missed entirely. But when the next 1D scan line revisits that DNA location, enHD will be ∼170 nm away and back in the high-friction mode, which should produce gaps in fluorescence of about 1-2 pixels on the kymograph (85 nm per pixel). Fluorescence gaps consistent in length and frequency with such phenomenon are readily apparent in the experimental 1D diffusive trajectories of enHD (e.g. see Fig. 4). Given the frequency and duration of such gaps, we deem it highly unlikely that they are caused by fluorophore blinking, which should be rare at the very low irradiance conditions of these experiments, and also much shorter lived (< 1ms). Fully resolving these events will require combined improvements in the optical and time resolutions of single-molecule fluorescence tracking methods. In the interim, we can conclude that the long diffusive trajectories of enHD do have the signatures expected for the hybrid scanning motion inferred from the analysis of the *D_1D_* variance.

We thus propose that enHD can navigate rugged DNA binding landscapes because it mimics the search strategy of a stochastic gradient descent algorithm (*66*). In this analogy, the high friction mode plays the role of the local gradient descent optimization steps, each consisting of ∼85 bp DNA sweeps performed at ∼1 bp/ms. The low-friction mode provides the stochastic sampling by enabling enHD to escape from a local minimum and redeploy to another location. The stochastic high-low-friction cycles of enHD thus emerge as an elegant solution to solving the speed vs. stability paradox (*8, 14, 67, 68*) for cases in which the DNA binding landscapes are energetically rugged.

### The high-friction mode: continuous sliding or mixed with hopping and gliding?

We should emphasize that even the high-friction mode alone is still many orders of magnitude faster than expected for a continuous promiscuous slide. This realization strongly suggests that enHD does not slide continuously while in the high-friction mode, but rather uses a mix of short-lived sliding runs connected by other nanoscale events such as hops and/or glides. Hopping is a process by which the protein briefly detaches from the DNA without escaping from the surrounding electric field and hops along its length, landing on a close location (*10, 11*). Gliding is a slightly different motion in which the protein engages in a series of ‘kiss and ride’ contacts with the phosphate backbone to move along the DNA with minimal friction (*69*). Either of these local modes has proven very difficult to resolve in single-molecule tracking experiments, likely due to general limitations in the temporal and spatial resolution of fluorescence imaging methods. On the other hand, there is mounting evidence that long 1D diffusive trajectories are indeed composed of a mix of sliding, hopping, and potentially gliding motions. Such a mix of motions at the nanoscale is consistent with the results from bulk kinetic experiments that probed DNA translocation of DNA repair proteins, and which indicate that continuous sliding may cover less than 10 bp (*70*). A recent high-resolution single-molecule study of the rotational component of 1D diffusion on DNA has also shown that the Lac repressor takes ∼40 bp to perform a full rotation around the DNA, indicating that it alternates between ∼20 bp hops and sliding runs every 0.5 ms (*71*). In this regard, it is interesting that the extrapolation of the enHD stepping rate (*D_1D_*) to ‘zero friction’ is at ∼1/4 of the slowdown from linear diffusion that is expected for pure rotational sliding (Fig. 6B). Propagation errors notwithstanding, this extrapolated rate is consistent with enHD covering ∼4 DNA helical turns, or 44 bp, per full rotation. Hence, the enHD high-friction mode is strikingly consistent with the 1D diffusion properties that have been inferred from experiments sensitive to nanometer scale motions on other proteins.

### The low-friction mode: 3D diffusion or long jumping?

A plausible interpretation of the low-friction mode is that it consists of brief 3D diffusion excursions in which enHD detaches from the DNA, moves to bulk to perform a short diffusion-collision search, and then lands back on the DNA to continue the scanning process at another location. However, the distances along the DNA that we estimate enHD travels while in the low-friction mode appear to be too long and fast to occur via 3D diffusion. Particularly, our results indicate that enHD takes 3-4 ms to travel an average of ∼173 nm in the low-friction mode. A freely diffusing enHD molecule (*D* ∼10^−6^ cm^2^⋅s^−1^) would only need ∼40 μs to diffuse 173 nm away from its starting position. But in the absence of directional bias, such distance would be traveled in any 3D direction, highly reducing the probability for landing on a ∼300 bp DNA target (per our position accuracy) located at such distance on either side of the mechanically extended DNA. As a way to estimate this probability, we calculated the ratio between twice the cylindrical capture volume of a 300 bp section of DNA, which accounts for the possibility of landing on either side of the DNA, and the volume of a sphere with 173 nm radius. This simple calculation does render a very small landing probability. In fact, even assuming a generous 3 nm radius for the capture cross-section of DNA (i.e. ∼2 nm around the DNA perimeter), the estimated landing probability is ∼10^−4^, which translates to an effective search time about 100-fold longer than the 3-4 ms of enHD’s low-friction bursts. Therefore, our experimental results suggest that enHD does not perform unbiased 3D diffusion during the low-friction bursts, but remains spatially correlated with the DNA as it diffuses along its length.

The low-friction bursts are thus more like long jumps on the DNA that connect high-friction scanning segments and occur in stochastic series until the protein manages to escape to bulk, ending the 1D diffusion trajectory. As discussed above, it seems that the only physical factor that could possibly maintain enHD’s contactless spatial correlation is the electrostatic attraction to the DNA backbone. But such electrostatic force may perhaps need to be farther reaching than the ∼1 nm currently estimated for the Debye length (*9*) to be able to maintain a correlated contactless motion over longitudinal displacements of 100 nanometers or more. A far-reaching DNA electrostatic field has been proposed by others as the reason behind the very long 1D diffusion trajectories observed for DBPs (*70*). The low-friction mode of enHD provides further evidence in support of a DNA-induced electrostatic field that extends over significantly longer distances. However, to sort this critical issue out an accurate experimental determination of the effective Debye length for protein-DNA interactions is needed.

### Functional implications for the control of gene expression programs in eukaryotes

The DNA scanning properties of enHD provide interesting functional insights for the control of transcription in eukaryotes. The DNA scanning needs of eukaryotic TFs are in principle far more demanding than just searching for one target site in one operon. Eukaryotic TFs often control multiple, even hundreds, of genes that can be distributed among multiple chromosome territories. This is particularly the case for master regulators or pioneer TFs, which are in charge of global gene expression programs. Individual genes can also depend on several regulatory elements, some of which are localized nearby acting as *cis*-elements (*72*) while others are placed at much longer distances operating in *trans* as enhancers (*43*). It appears evident that the simultaneous operation in a multitude of chromosome territories may need a specialized mechanism for tracking target genes and regulatory elements that eliminates the need for continuously scanning the full genome. One such tracking mechanism has been recently proposed, which combines the recent discovery of promiscuous DNA recognition with the repetitive clusters of imperfect cognate motifs that are found in the regulatory regions of eukaryotes (*39*). In this mechanism the clusters of imperfect motifs operate as a transcription antenna that draws molecules of a promiscuous TF by engaging them in myriads of local, mid-affinity binding events. Bioinformatic analysis has suggested that the regulatory regions of the 203 genes known to be under direct enHD control may provide sufficient promiscuous binding power to dynamically capture most of the 30,000 enHD molecules that are present in a living cell, leaving just a few free in bulk solution (*39*). Transcription antennas can thus potentially solve the tracking problem while providing a key functional role for the degenerate sequence architectures of the regulatory regions in eukaryotes.

Nevertheless, once the TF is circumscribed to a given chromosome territory it must still scan large segments of DNA to perform its function. This regional scanning process might also be quite different from the search for one DNA target site since it involves finding and/or recruiting cofactors, chromatin remodelers, or even additional copies of the same TF in/to DNA regions that are dynamically packed and unpacked. Hence the same rugged DNA binding landscapes that serve to attract promiscuous TFs could impair their ability to efficiently scan through the DNA region of interest. For example, according to our calculations, enHD would take one hour to scan just a 1 kbp piece of the λ-phage DNA using a continuous sliding motion. The implication is that promiscuous TFs such as enHD, and presumably any TF with a very short cognate motif, need an alternative mechanism for DNA scanning. In addition, many of these TFs forego a DNA clamping interface so they can target DNA on nucleosomes, which imposes further demands for achieving extensive DNA scanning. In this regard, our results provide direct demonstration that such TFs can indeed scan DNA fast and extensively. We find that enHD moves along rugged DNA binding landscapes about ten million fold faster than it would via a continuous sliding motion. We also find that the key for scanning rugged DNA landscapes is a hybrid mechanism that alternates stochastically between high and low friction modes. Interestingly, even though we have studied the DNA scanning of enHD only in naked mechanically extended DNA, we note that the mechanism it uses under such conditions may still shed some light on how pioneer TFs scan active and silent DNA regions *in vivo*.

For instance, enHD covers about 27.5 nm (85 bp) per run of its high-friction mode, which is when it truly scans the DNA sequence. These traveled distances happen to be very close to the perimeter of a single nucleosome (34 nm), or to the amount of wrapped DNA that is accessible on the nucleosome surface (∼61% of 147 bp). Although the binding to nucleosomes of enHD has not been studied directly yet, its DNA binding properties are identical to those of other homeodomains known to target nucleosome-wrapped DNA all around the nucleosome perimeter (*38*). Single-molecule studies of the yeast TFs Reb1 and Cbf1 have found that these proteins bind nucleosomes with the same affinity than naked DNA, albeit with slower overall dynamics (*73*). If we assume a similar behavior for enHD, the high-friction mode we observe in naked extended DNA could perhaps reveal a mechanism optimized for scanning DNA in single nucleosome units. In this context, a cluster of imperfect cognate motifs would significantly increase the time the TF remains associated with the cluster-bearing nucleosome, potentially facilitating the TF’s ability to find or recruit other partners on/to the site. A cluster of imperfect motifs could also promote local swarming of TF molecules, which could then collectively induce changes in DNA packing: *i*) on naked DNA the TF swarm could dynamically block the reassembly of nucleosomes to keep the region transcriptionally active; *ii*) whereas on nucleosomes, a local TF swarm could either stabilize the DNA in the packed configuration, or potentially cause histone eviction if binding were to perturb the nucleosome structure (*37*).

The low friction mode of enHD provides a simple escape mechanism, which might be of particular importance for scanning chromatin without getting trapped on the highly sticky binding dynamics that are likely to ensue with clustered-bearing nucleosomes. Using such an escape mechanism the TF could thus leap from one clustered region to the next one within the regulatory locus of interest. EnHD long jumps ∼173 nm in the low-friction mode. In naked extended DNA, these long jumps correspond to ∼520 bp, but they would presumably result in longer sequence displacements if they occurred in similar fashion on nucleosome packed DNA. From a functional standpoint, it has been noted that the clusters of imperfect motifs in the regulatory regions of higher-order organisms are often organized as archipelagos (*74*). In this regard, we note that the distance displacements that occur during the low-friction mode of enHD are consistent with the sequence spacings found between the islands of clustered-motifs of regulatory elements. It is thus tempting to speculate that the low-friction mode perhaps provides an evolved strategy for efficiently shuttling among such islands.

These considerations lead us to propose that the complex sequence architecture of eukaryotic regulatory elements may in fact represent a sophisticated code that defines the specific level of transcriptional activation from the number of pioneer TF molecules that can be dynamically recruited to the locus of interest. Patterns of recruitment could be encoded in the size, number, spacing, and exact positioning of the clusters of imperfect cognate motifs for relevant TFs. Such a programmable TF recruiting system would be able to add another dimension to the classical feedback control mechanism based on the up- or downregulation of the cellular TF concentration. These ideas obviously need to be further investigated via carefully designed single-molecule and *in vivo* experiments. On the other hand, given the paradigmatic DNA binding and functional properties of Engrailed, it is quite possible that the DNA scanning process hereby outlined for enHD is commonly used by other eukaryotic factors that function as pioneer TFs.

## Materials and Methods

### Protein expression, purification, and labeling

The enHD protein used in this study is identical to the one we used to characterize binding promiscuity (*39*), and corresponds to the sequence: MAEKRPRTAFSSEQLARLKREFNENRYLTERRRQQLSSELGLNEAQIKIWFQNKR AKIKKSTC We also studied the binding properties of the Q50K mutant of enHD (*59*). This variant was produced by site-directed mutagenesis of the gene encoding for the wild-type sequence. The protein was expressed and purified from *E.coli* BL21 star DE3 (ThermoFisher scientific) cells using a previously published protocol (*39*). 200 mL LB broth with 100 μg/mL Ampicillin was inoculated with a colony of freshly transformed *E. coli* Bl21 Star DE3 cells with pBAT 4 plasmid containing the gene encoding EnHD, and grown to an O.D. of 1 at 37°C, 250 RPM in an incubator shaker. The primary culture was transferred into 2-liter terrific broth and induced with 1 mM IPTG (isopropyl-β-D-thiogalacto-pyranoside) at an O.D. of 1.2 and grown overnight at 37°C, 250 rpm shaking. Cells were harvested by centrifuging at 8,000 rpm for 20 minutes, and the pellet was lysed by incubating with the bacterial protein extraction reagent (B-PER^TM^, ThermoFisher Scientific) on a rocker for 30 minutes at RT. Cell debris was removed by centrifuging at 30,000 rpm for 30 minutes in an ultra-centrifuge. The supernatant was loaded on to a SP Sepharose column (HiTrap SP column, GE healthcare) and washed with 50 mM Phosphate buffer with 100 mM NaCl, pH 6.5 and the protein was eluted with a gradient of 50 mM Phosphate buffer with 1 M NaCl pH 6.5. The elution fractions containing the protein were pooled and further purified using reverse phase chromatography using a C4 column (Higgins Analytical, Mountain View, CA, USA) with 5% Acetonitrile and 0.1 % trifluoro acetic acid as mobile phase A and 95% Acetonitrile and 0.1% trifluoro acetic acid as mobile phase B. The fractions containing the enHD were pooled and flash frozen and lyophilized. The C-terminal cysteine residue was labeled with Alexa 488 C5-maleimide (ThermoFisher Scientific). The labeling was performed in 50 mM phosphate buffer at pH 7 with 100 mM NaCl and 8 M Urea. The protein was incubated with 2-fold molar excess of Alexa 488 C5 Maleimide and incubated at room temperature for 2 hours. Excess unreacted dye was separated from the protein by cation exchange chromatography using SP Sepharose column, after washing away the free unbound, unreacted dye. The protein was eluted with 1 M NaCl and the fractions containing the labeled protein were pooled and flash frozen and stored at −80 °C.

### Tracking of Alexa 488-labeled EnHD binding on lambda DNA

All the experiments of enHD binding to lambda DNA were performed on commercially available dual beam optical trap coupled to confocal microscopy (C-Trap, Lumicks). All the experiments were performed in 20 mM Tris buffer at pH 7.5 with NaCl concentrations ranging from 25 mM to 100 mM. The buffer also contained photo protection reagent comprising of 100 μg/mL Glucose oxidase, 20 μg/mL catalase, 5 mg/mL Glucose and 1mM Trolox. 0.05% v/v Tween 20 was added in all buffers to prevent the labeled protein from sticking to surfaces of the tubes and flow cell. Biotinylated double stranded lambda DNA was tethered in between two streptavidin coated polystyrene beads of 3.11 μm diameter. The binding of Alexa 488 labeled enHD to the lambda DNA was probed by performing fluorescence line scans along the length of the DNA using a confocal microscope. The pixel size used in the fluorescence scan is 200 nm and each pixel was imaged for 50 μs. The fluorescence line scans taken at every ten ms was aligned in series to make a kymograph whose x-axis represents time and y-axis represents position along the lambda phage DNA.

### Stochastic binding simulations

Stochastic binding simulations were performed in MATLAB. A total of 25000 1D diffusion trajectories were simulated with low friction and high friction mode of 1D diffusion, and the number of switches between the two modes was changed to different values (1, 2, 3, 6, 10). The diffusion coefficient used for low friction mode is the translational diffusion of enHD calculated using Stokes-Einstein equation with a low friction of 0.5 RT to account for the minimum interaction the protein has with DNA. For the diffusion coefficient for high friction mode, we used the equation described earlier for sliding (Schurr, 1979), with a friction term calculated from the average binding energy of enHD on lambda DNA calculated from our previously published statistical mechanical model (Castellanos et al., 2020).

### Data analysis

The kymographs obtained from the C-trap instrument were analyzed using a custom algorithm written in python (*75*). This algorithm identifies binding events by grey scale dilation of the original image following by targeting of the pixels that remain unchanged. The specific location of the fluorophore (protein) is then determined with sub pixel accuracy by a refinement step that calculates the offset of a brightness-weighted centroid around the vicinity of each pixel. The subsequent motion of the protein is tracked by connecting pixels in the adjacent frames using the greedy algorithm (*75*).

The position and time information of each binding trajectory derived from the python script were further analyzed in MATLAB to determine the dwell time on the DNA molecule, total distance traveled along the DNA, and the diffusion coefficient of each trajectory using custom-built code. The average dwell time is calculated from the distribution of dwell times obtained from the individual 1D trajectories. The total distance traveled per 1D diffusive event is obtained by summing the distance displacements between time points over the entire trajectory. The diffusion coefficient for each trajectory is calculated using the formula:

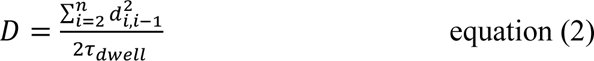

where *n* is the total number of time points along the trajectory, *d_i,i_*_−1_ is the displacement of the protein between times points *i-1* and *i*, and τ*_dwell_* is the duration of the 1D trajectory. The *D_1D_* in cm^2^/s is converted into stepping rate using the formula:

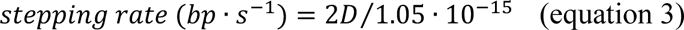

### Determination of position accuracy of the dual-beam optical tweezers coupled to confocal fluorescence microscope

The position accuracy along the DNA of the dual beam optical tweezers coupled to confocal fluorescence microscope (C-Trap, Lumicks) was determined by imaging a λ-DNA molecule modified to incorporate one copy of the Atto488 fluorophore at the 33,786 bp position (Lumicks). The Atto488 λ-DNA molecule was tethered to two streptavidin coated polystyrene beads, mechanically controlled by the dual traps and imaged with the correlative scanning confocal fluorescence microscope.

The kymographs were acquired at a pixel size of 100 nm with 50 µs photon collection per pixel. The fluorescence intensity profile along the DNA position was obtained by summing up the counts as a function of time at each pixel from the kymograph. The specific position of the fluorophore along the DNA was determined by point spread function analysis of each kymograph line scan using a code developed earlier (*76*). The distance axis of the kymographs was converted to base pairs on the λ-DNA by locating the fluorescence intensity peaks originating for the flanking beads to delimit the DNA ends and using the 16.5 µm contour length of the λ-DNA. The location of the fluorophore determined using point spread function analysis of the kymograph line scans, which resulted on a distribution with a standard deviation of 53.5 nm or 156 bp, indicating that the position accuracy on the DNA is 312 bp at 95% confidence (Fig S1).

## Author contributions

Conceptualization: VM

Methodology: RG, MS, SN, VM

Investigation: RG, MS

Analysis: RG, VM

Supervision: VM

Writing: RG, VM

## Data and materials availability

All data, code, and materials used in the analyses are available upon request to the corresponding author.

## Supplementary Materials

**This PDF file includes:**

Figs. S1 to S6

