## Supplemental Materials for "How to Scan DNA using Promiscuous Recognition and No Sliding Clamp: A Model for Pioneer Transcription Factors"

**This PDF file includes:**

Figs. S1 to S6

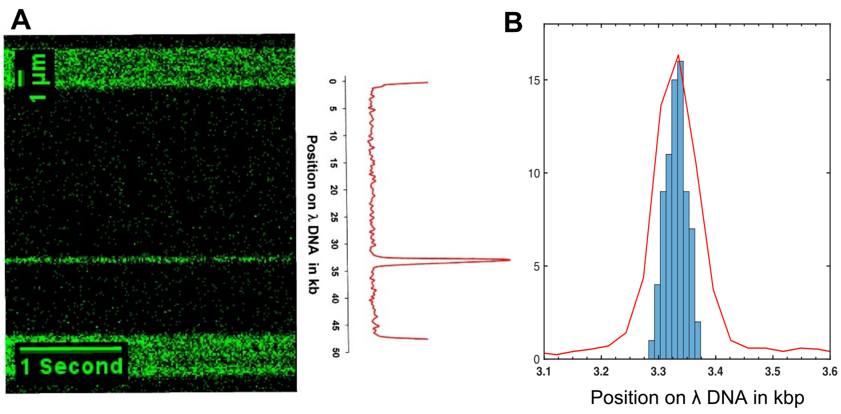


**Figure S1**. Experimental determination of the position accuracy on the DNA of the dual beam optical trap coupled to confocal fluorescence microscope system. (**A**) Kymograph of the λ-DNA labeled with Atto488 label at position 33,786 bp tethered between two ~2.1 μm diameter polystyrene beads. The position along the λ-DNA is converted to kbp using the known contour length of the λ-DNA (16.5 μm). The fluorescence intensity profile integrated along the time axis is shown on the right in red showing the location of the fluorophore at ~33.7 kb. (**B**) Histogram showing the distribution of the fluorophore's position obtained from the point spread function analysis performed with the tracking algorithm on all of the line scans of the kymograph from panel A. The position of the fluorophore is determined with a standard deviation of 53.8 nm or 156 bp, indicating a position accuracy of ~312 bp at 95% confidence.

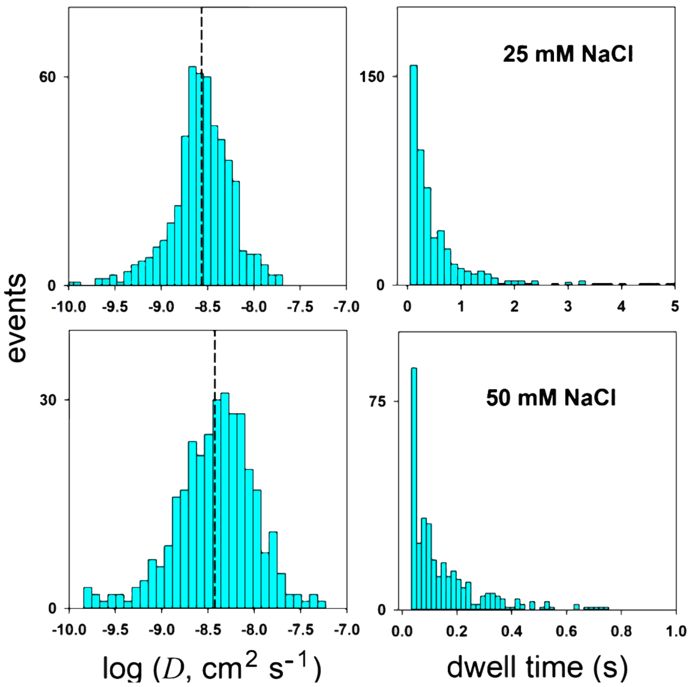


**Figure S2**. Histograms showing the distribution of 1D diffusion coefficients in $\log_{10}$ units (left panels) and dwell times (right panels) from individual diffusing trajectories of the wild-type enHD measured at 25 mM NaCl (top), 50 mM NaCl (bottom). The mean value for each distribution is shown with a dashed line.

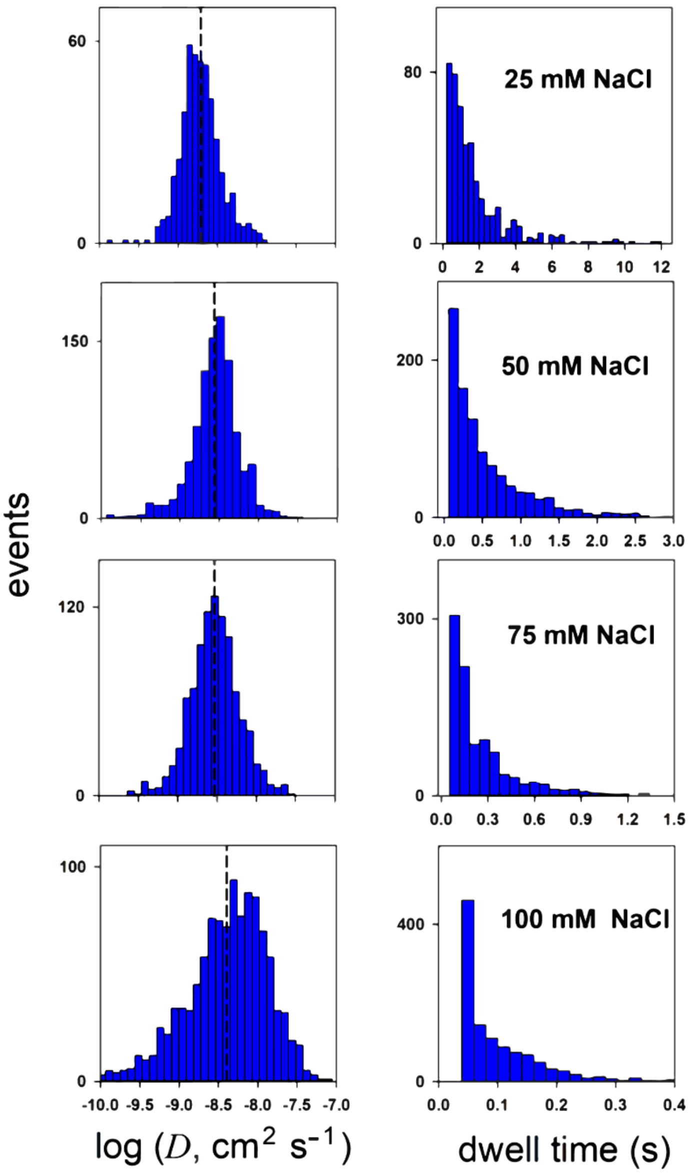


**Figure S3**. Histograms showing the distribution of 1D diffusion coefficients in $\log_{10}$ units (left panels) and dwell times (right panels) from individual diffusing trajectories of the Q50K variant of enHD measured at different NaCl concentrations. The diffusion coefficients increased with increase in salt concentration and the distribution became broader with increase in salt. The mean value for each distribution is shown with a dashed line. The dwell times followed an approximately exponential distribution at each salt concentration with the exception of the highest salt concentration.

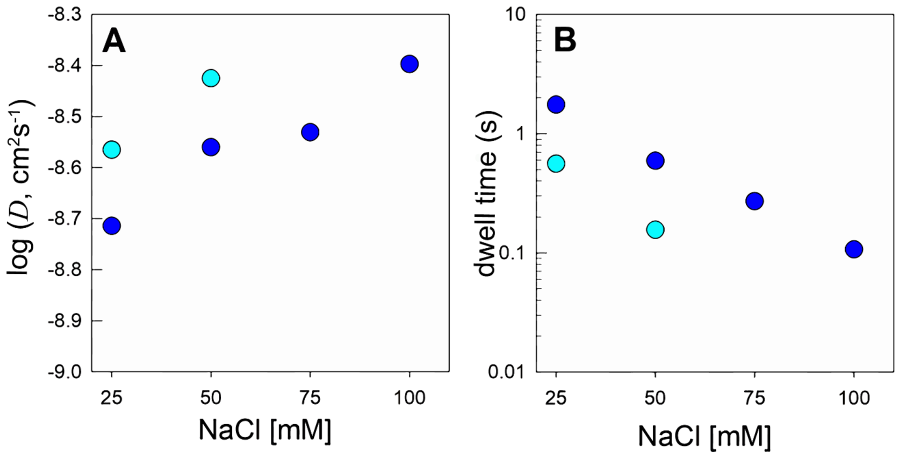


**Figure S4.** Effects of the salt (NaCl) concentration on the scanning of the λ-DNA by enHD wild-type (cyan) and Q50K (blue) variants. The circles represent the mean measured values for th1D diffusion coefficient in $\log_{10}$ units (**A**) and dwell time (**B**) obtained from the histograms of Figs. S1 and S2.

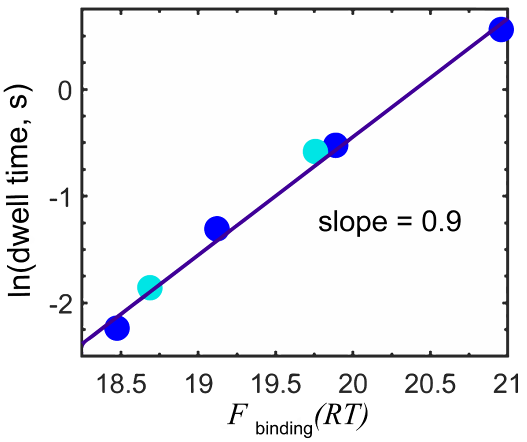


**Figure S5**. Experimental mean dwell time (in natural log units) versus the average free energy of binding of enHD to the λ-DNA from the statistical mechanical model. Data for wild-type and Q50K variant of enHD are shown in cyan and blue circles, respectively.

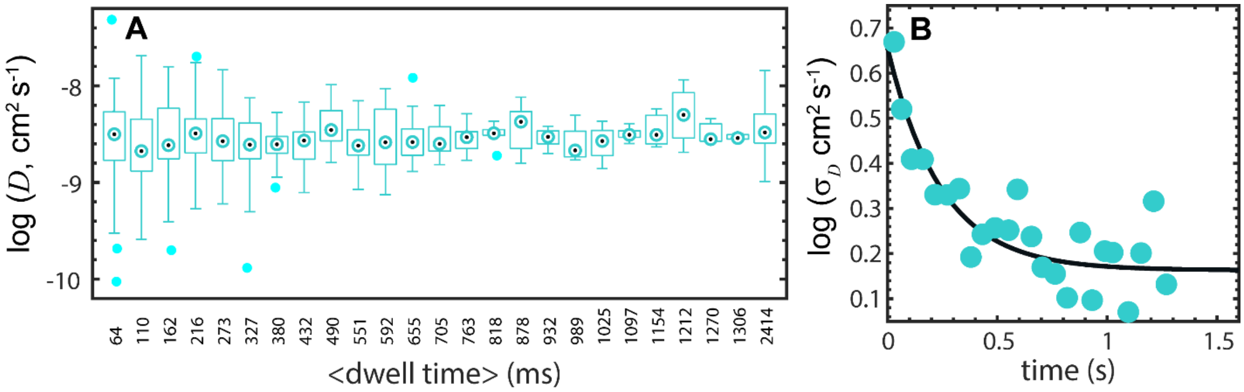


**Figure S6**. (**A**) Box plot showing the variation in overall trajectory $D_{1D}$ as a function of the duration of the scanning trajectory on the λ-DNA for the wild-type enHD. Whiskers show the end points, box edges the lower-upper quartiles, and dotted circles the bin medians. Cyan circles are outliers. (**B**) The standard deviation for the data within each bin of panel **A** in log10 units versus the characteristic binned time. All of the data for trajectories longer than 1200 ms were combined into a single group to minimize statistical errors due to the small numbers.
